# Investigating the attentional focus to workplace-related soundscapes in a complex audio-visual-motor task using EEG

**DOI:** 10.1101/2022.10.06.511066

**Authors:** Marc Rosenkranz, Timur Cetin, Verena Uslar, Martin G. Bleichner

## Abstract

In demanding work situations (e.g., during a surgery) the processing of complex soundscapes varies over time and can be a burden for medical personnel. Here we study, using mobile electroencephalography (EEG), how humans process workplace-related soundscapes while performing a complex audio-visual-motor task (3D Tetris). Specifically, we wanted to know how the attentional focus changes the processing of the soundscape as a whole. Participants played a game of 3D Tetris in which they had to use both hands to control falling blocks. At the same time, participants listened to a complex soundscape, similar to what is found in an operating room (i.e., sound of machinery, people talking in the background, alarm sounds, and instructions). In this within-subject design, participants had to react to instructions (e.g., “place the next Block in the upper left corner”) and to sounds depending on the experimental condition, either to a specific alarm sound originating from a fixed location or to a beep sound that originated from varying locations. Attention to the alarm reflected a narrow attentional focus, as it was easy to detect and most of the soundscape could be ignored. Attention to the beep reflected a wide attentional focus, as it required the participants to monitor multiple different sound streams. Results show the robustness of the N1 and P3 event related potential response during this dynamic task with a complex auditory soundscape. Furthermore, we used temporal response functions to study auditory processing to the whole soundscape. This work is a step towards studying workplace-related sound processing in the operating room using mobile EEG.

## 1 Introduction

Auditory attention, i.e., focusing on relevant and ignoring irrelevant sounds, is a fundamental skill at workplaces with both, a high responsibility and a soundscape containing a variety of sounds. Surgery staff, for example, performs a highly complex task while being exposed to conversations, machine and tool sounds, and background music. This soundscape accumulates to sound pressure levels regularly exceeding 50dB(A) (Baltin et al., 2020; Engelmann, Neis, Kirschbaum, Grote, & Ure, 2014; Hasfeldt, Laerkner, & Birkelund, 2010; Tsiou, Efthymiatos, & Katostaras, 2008). Surgery staff reported that the soundscape can become a burden (Healey, Primus, & Koutantji, 2007; Pleban, Radosz, Kryst, & Surgiewicz, 2021; Tsiou et al., 2008; van Harten, Gooszen, Koksma, Niessen, & Abma, 2021), and increases surgical complication rates (Engelmann et al., 2014; Kurmann et al., 2011). Interestingly, the focus of attention to sounds and their interpretation changes throughout a surgery. For example, conversations are sometimes perceived as disturbing as the staff member must concentrate on the task, while other times they are attended and even encouraged (van Harten et al., 2021). In the former case auditory attention is narrowed to task-relevant sounds (e.g., instructions) and tries to suppress irrelevant sounds. In the latter case the attentional focus is wide, i.e., switches between multiple sound sources. Our goal was to study a narrow compared to a wide focus to better understand auditory attention in a complex and multi-sensory environment.

Electroencephalography (EEG) can be used to measure auditory attention continuously, objectively and without the interruption of a person. Mobile EEG systems allow to study the brain in a working environment rather than in the lab (Hölle, Meekes, & Bleichner, 2021; Wascher, Heppner, & Hoffmann, 2014; Wascher et al., 2021) and has already been used to assess performance during laparoscopic training (Shafiei et al., 2021; Suárez et al., 2022; Thomaschewski et al., 2021). Thus, with EEG we want to study auditory attention in the operating room and understand when sounds become a burden.

When transitioning from the lab to the operating room we must consider that our expectation about auditory attention is mainly derived from highly controlled studies (Gramann et al., 2021). In order to generalize lab findings to more complex environments we have to increase the environmental complexity. One approach to increase complexity is naturalistic laboratory research which provides a balance between stimulus control and ecological validity (Matusz, Dikker, Huth, & Perrodin, 2019). We decided to develop a complex and dynamic, audio-visual-motor task while maintaining experimental control over stimuli. Thereby, EEG responses related to auditory attention can be studied in a complex environment.

We first operationalized the soundscape of an operating room into five stimulus categories: a continuous background stream, as well as, task relevant and irrelevant sound and speech stimuli (Engelmann et al., 2014; Hasfeldt et al., 2010). The background stream represents sounds originating from running machines, ventilation, and people moving around. Task relevant speech represents exchanges about the surgery, as well as, instructions. Task irrelevant speech represents private conversations. Task relevant sounds represent, e.g., alarm sounds and feedback from instruments. Task irrelevant sounds represent, e.g. phone ringing or sounds from monitors.

We then combined our operationalization of the soundscape with a visual-motor task, namely the computer game Tetris. The game requires the use of both hands to navigate blocks. For the continuous background stream we chose a hospital soundscape. For task relevant speech participants received instructions within the game. For task irrelevant speech a conversation unrelated to the game was presented. The task relevant sound changed between two conditions. For task irrelevant sounds monitor sounds from a surgery machine were presented.

Lastly, we manipulated the attentional focus of the participants by changing the task relevant sound while keeping the complexity of the soundscape constant. In a narrow attentional focus condition (from here on narrow condition) participants had to attend to an alarm sound (from here on the alarm). This sound originated from a specific location, i.e., was easy to detect. The rest of the soundscape (except the task relevant speech) could be ignored. In a wide attentional focus condition (from here on wide condition) we implicitly direct the participants attention towards all sound streams by instructing participants to attend to a sound that was embedded in any of the five streams. We refer to this sound as the beep, as it served the purpose of manipulating participants attention but was generally unrelated to the operation soundscape.

The aim of our study was twofold. First, we investigated well-known EEG responses, namely event-related potentials (ERPs) and temporal response functions (TRFs) in a dynamic task with a complex soundscape using a mobile EEG setup. Second, we investigated neural processing when the auditory attentional focus was narrow compared to wide.We used ERPs to study responses to distinct stimuli, i.e., relevant and irrelevant sounds, and focused on two components, the N1 and P3 response: The N1 response is an early negative deflection related to auditory processes and modulated by attention (Hansen & Hillyard, 1980; Hillyard, Hink, Schwent, & Picton, 1973; Luck, 2014; Näätänen & Picton, 1987; Picton & Hillyard, 1974). For our first hypothesis, we expected a larger N1 response for irrelevant sounds (i.e., non-target sounds in both conditions) in the wide condition than in the narrow condition. Attention to the beep, which was integrated into other sounds, should lead to a stronger processing of the whole soundscape. Therefore, we expected also a stronger processing of the irrelevant sounds.

The P3 response is a late positive deflection in response to target sounds (from here on targets). As this response is absent in non-targets it thereby marks attentional processes (Luck, 2014; Polich, 2007). For our second and third hypotheses we expected a P3 response to the target of the respective conditions. The alarm was the target in the narrow condition, thus we expected a larger response in this condition compared to the wide condition. The beep was the target in the wide condition, thus we expected a larger response in this condition compared to the narrow condition.

We used TRFs to investigate processing of the soundscape as a whole. TRFs are the result of correlating a continuous EEG signal with a continuous audio signal (Crosse, Di Liberto, Bednar, & Lalor, 2016). The correlation (i.e., response) is larger for attended compared to unattended signals (Mirkovic, Debener, Jaeger, & De Vos, 2015; O’Sullivan et al., 2015). For our fourth hypothesis we expected that the TRF is larger in the wide compared to the narrow condition, as the beep should direct attention towards the whole sound environment.

## 2 Method

This study was registered prior to any human observation of the data (https://osf.io/sgvk6). Deviations from our preregistration are described in the supplementary material. We provided the experiment, as well as the code and data to reproduce the statistical analyses and figures here: Rosenkranz and Bleichner (2022).

### 2.1 Participants

Twenty-two participants (age range: 20-30 years; female: 16) were recruited through an online announcement on the University board. We based the sample size on previous studies showing P3 effects in natrualistic settings (e.g., Hölle, Meekes, & Bleichner, 2021; Protzak & Gramann, 2018; Scanlon, Townsend, Cormier, Kuziek, & Mathewson, 2019) due to the exploratory approach of this study. All participants signed prior to the experiment informed consent approved by the medical ethics committee of the University of Oldenburg and received monetary reimbursement. Eligibility criteria included: Normal hearing (self reported), normal or corrected vision, no psychological or neurological condition, righthandedness, and compliance with current COVID-related hygiene regulations (e.g., this could include proof of vaccination).

Two participants were excluded from the final analysis. One participant showed high impedance (*>*100 kΩ) at the end of the experiment and overall poor data quality. One participant had a very low hit rate which indicates that this participant did not follow task instructions. I.e. 20 participants were included in the final analysis.

### 2.2 Paradigm

Participants performed a complex audio-visual-motor task - an adapted 3D Tetris game. The basis for the game was developed by Kalarus (2021) and we changed it to our needs. Below is a short description of the paradigm. For a detailed description of the game and generation of the auditory stimulus material see Supplementary Method.

Participants had to play a 3D Tetris game while reacting to different sounds and instructions. In 3D Tetris one is presented with a three-dimensional space in which differently shaped, three-dimensional blocks are placed. The falling blocks must be placed in such a way that they form a layer. When a layer is complete, the layer disappears. Participants controlled the rotation of the blocks with the left and position with the right hand. The participants goal was to place blocks to remove as many layers as possible to receive points. Unlike the classic Tetris game, participants could not loose when the blocks were stacked too high. Instead the game restarted at the bottom layer to allow for a continuous game-play. When that happened, participants lost points.

Furthermore, participant were listening to a soundscape. The soundscape included one continuous background sound, and five discrete stimuli (see Figure 1B): The background sound consisted of hospital sounds containing e.g., air conditioning and people moving around and was presented from both sides. A task irrelevant speech of two people talking in the background originated to the left behind the participant (−135°). Two irrelevant sounds were presented from the left side (−90°). Participants also received from time to time instructions from the front (0°) where to place the next block. Furthermore, an alarm was presented from the right (45°) and a beep could occur from the same direction as other stimuli. All sounds were spatially separated using the Head Related Impulse function (Kayser et al., 2009).

**Figure 1:**
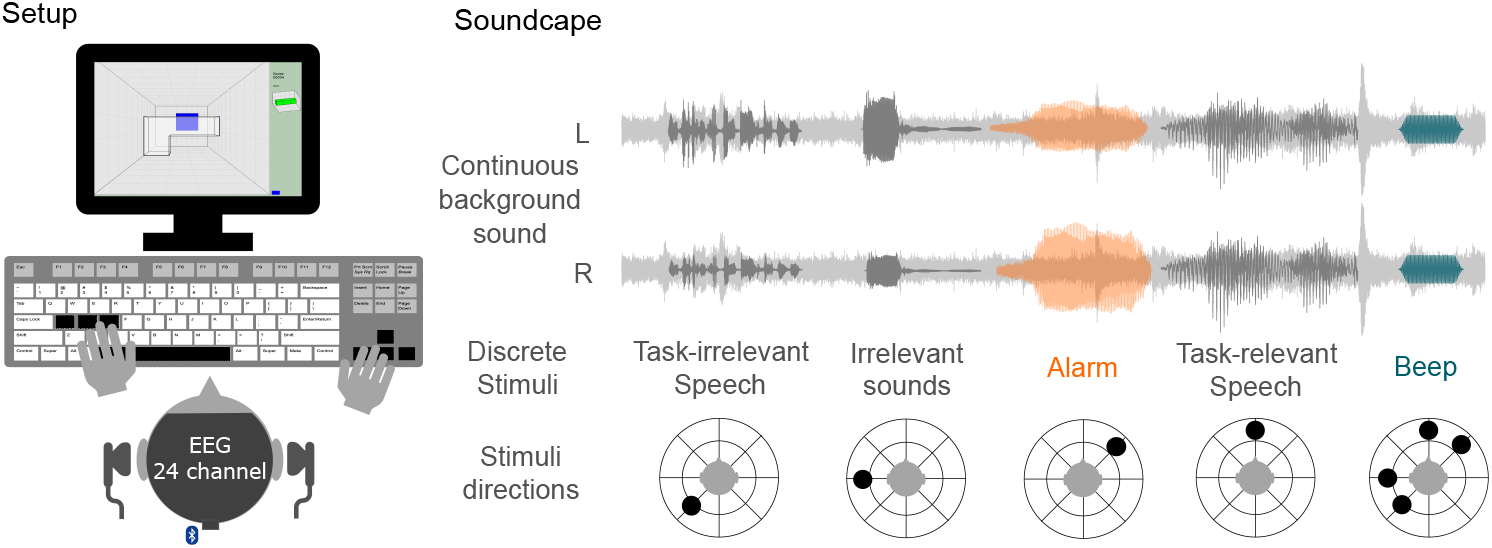
**A)** Experimental Setup: Participants played 3D Tetris (with the left hand participant controlled the rotation on a block, with the right hand the position of a block). The soundscape was presented via headphones. EEG was recorded using a 24 channel mobile EEG setup. **B)** Soundscape: A continuous background sound was presented throughout the task. Discrete stimuli were subsequently presented. The alarm was the target in the narrow condition, while the beep was the target in the wide condition. The alarm was presented from one direction, the beep was presented from any direction as the other sounds. If participants detected a target, they should press the space bar.

For the auditory task, participants should always attended to the task relevant speech, which instructed participants to place the next block in one of the four corners of the Tetris layer. Furthermore, participants played the game twice (a game lasted approximately 18 minutes) and received a different instruction for each condition. Note, that the soundscape was conceptually the same for both conditions. In the narrow condition, participants were instructed to additionally attend to the alarm, i.e., participants had to attend to the task relevant speech and the alarm. The alarm was long, had a high intensity, and was presented always from the same direction, thus the rest of the soundscape could be ignored to detect it. In the wide condition, participants were instructed to attend to the beep, i.e., participants had to attend to the task relevant speech and the beep. The beep was short, had a low intensity, and was integrated into other stimuli, thus the whole soundscape had to be monitored to detect it. To summarize, the difference between the two conditions were the instruction which target should be attended to. The target of the narrow and wide condition were the alarm and the beep, respectively. If participants detected a target they should press the space bar. Hitting a target, as well as, following the speech instructions granted points while misses and not following instructions subtracted points.

All discrete auditory stimuli were initially presented 48 time in a random order. However, the response to the beep overlapped with the response to the alarm and irrelevant sounds when it was integrated into them. Therefore, we added all overlapping sounds again to derive at 48 non-overlapping sounds. Note, that only responses to non-overlapping sounds were used in the ERP-analyses.

To get acquainted to the game, participants received written instructions for the game. Then, a general training without auditory stimuli and a training for the relevant speech instructions was performed. Before each condition, participants performed also a condition specific training in which they received feedback whether they correctly detected the target (see Supplementary Method for a detailed description of the training games). EEG was not recorded during the training games. Before the game of each condition started, resting EEG was recorded, by instructing participants to first focus a fixation cross and then close their eyes for one minute each. Furthermore, two questionnaires were administered, which are not considered in the current study: At the beginning of the experiment participants filled out a noise sensitivity questionnaire (NoiSeQ; Schutte, Marks, Wenning, & Griefahn, 2007) and after each condition a workload questionnaire (NASA-TLX; Hart & Staveland, 1988).

### 2.3 Data acquisition

Participants were asked to wash their hair at the day of recording. EEG data was recorded using a wireless 24-channel amplifier (SMARTING, mBrainTrain, Belgrade, Serbia) attached to the back of the EEG cap (EasyCap GmbH, Hersching, Germany) with Ag/AgCl passive electrodes (see Supplementary Figure 3 for the channel layout) and the reference and ground electrode at position Fz and AFz, respectively. The data was recorded using a sampling rate of 500 Hz, and transmitted via Bluetooth from the amplifier to a Bluetooth dongle (BlueSoleil) that was plugged in a computer (Dell Optiplex 5070).

After fitting the cap, the skin was cleaned using 70% alcohol. To increase skin conductance between the scalp and electrodes abrasive gel (Abralyt HiCl, Easycap GmbH, Germany) was used. Impedance were kept below 20 kΩ at the beginning of the recording. Recording took place in a quite and electrically shielded room. Participants were seated in front of a screen (Samsung, SyncMaster P2470) and keyboard (Dell, KB 1421). Auditory stimuli were presented using Psychtoolbox 3 (v3.0.17, Kleiner et al., 2007). For each stimulus type, a sound marker was generated using the lab streaming layer library ^1^. A key capture software ^2^ was used to record which key was pressed on the keyboard and an audio capture software (this is used as input for the TRF analysis, see below) ^3^ was used to record the presented audio with a sampling rate of 44100 Hz. To synchronize all data streams, the transmitted EEG data, sound marker, keyboard marker, and computer audio was collected in the Lab Recorder software ^4^ based on the Lab Streaming Layer and saved as .xdf files. The same computer was used for data recording and experiment presentation.

### 2.4 Preprocessing

The EEG was analysed using EEGLAB (v2021.0, Delorme & Makeig, 2004) in MATLAB R2020b (The MathWorks, Natick, MA, United States).

For each participant and condition, the continuous data was filtered with Hamming windowed FIR filter using the EEGLAB default seetings: (1) high-pass: passband edge = 0.1 Hz^5^; (2) low-pass: passband edge = 30 Hz^6^. The filtered data was re-sampled to 250 Hz. Bad channels were visually inspected and removed from both conditions. Afterwards, the data was cleaned from artifacts using infomax independent component analysis (ICA) and rejected channel were interpolated.

For the ICA, a copy of the preprocessed data was high-pass filtered (passband edge = 1 Hz^7^) and cut into consecutive epochs of one second. Epochs with a global or local threshold of 2 standard deviations were automatically rejected. ICA decomposition was applied on the remaining epochs of both conditions. The resulting components were back-projected on the original preprocessed, but uncleaned data of each condition. Components related to eye-blinks, eye-movement, heart rate, and muscle movement were identified and removed using the EEGLAB toolbox ICLabel (Pion-Tonachini, Kreutz-Delgado, & Makeig, 2019) with a threshold of .9. On average, 2.8 (±1.32) components were rejected. Afterwards, previously rejected channels were interpolated using spherical interpolation. Then, channel were re-referenced to the linked mastoids (TP9/TP10). For all auditory stimuli, a constant delay of 19 ms between the sound marker and sound presentation was taken into account.

#### 2.4.1 ERP analysis

ERP analyses were was performed for the alarm, beep, and the two irrelevant sounds. For each of the sounds, epochs from −200 to 800 ms with respect to the stimulus onset were generated and a baseline correction from −200 to 0 ms prior to stimulus onset performed. Epochs with a global or local threshold of 3 standard deviations were automatically rejected. For targets (i.e. the alarm or beep) only hit trials were included in the analysis. A hit was defined as any space bar press within three seconds after a target.

We calculated ERP amplitudes averaged over time based on individual time-windows. Our ERP analyses focused on the N1 and P3 component. The analyses of the two components were identical, except for the selection of channel and time-window. For each participant, an average response was calculated from the two conditions and selected channels. For the N1, we selected channel Fz, FC1, FC2, Cz, C3, C4 and for the P3, we selected channel Pz, P3, P4, CPz, CP1, CP2 (see Supplementary Figure 3). The average response was used to find the component peaks of each participant. For the N1, we searched for a negative deflection between 50 and 150 ms following stimulus onset. For the P3, we searched for a positive deflection between 300 and 400 ms following stimulus onset. Following peak detection, the component time-window was determined. For this, a time-window of ±25 ms and ±50 ms around the N1 and P3 peak was taken, respectively. Lastly, to derive at trial-level data, for each participant, condition, selected channel, and trial, the mean amplitude over the individual time-window was calculated.

#### 2.4.2 TRF analysis

For the analysis of the soundscape as a whole the mTRF toolbox (Crosse et al., 2016) was used. Therefore, the recorded audio was preprocessed as follows: First, the absolute of the hilbert transform of the audio was low-pass filtered at 30 Hz^8^ and resampled to 250 Hz. Second, we were interested in the response to the whole soundscape irrespective of the targets, therefore, the alarm and beep were excluded from the EEG and audio data of both conditions. Data from the onset of the alarm and beep was excluded up to one second after the onset, creating epochs of unequal length. The total length of data was unequal between conditions, therefore we excluded epochs until the total length difference between conditions for each participant was less than one minute. Third, EEG data was multiplied by factor .0313 for normalization (as suggested in the provided code by Crosse et al. (2016)). Finally, a forward model was trained on the epoched EEG data and audio data using the function *mTRFtrain*. Time lags were calculated from −200 to 800ms and a lambda of 0.1 was be used.

The TRF usually reveals classic ERP peaks known from the auditory processing literature (Crosse et al., 2016; Jaeger, Mirkovic, Bleichner, & Debener, 2020; Mirkovic, Debener, Schmidt, Jaeger, & Neher, 2019). Based on pilot data from three participants (not included in the final analyses), we expected these peaks approximately around 100, 200, and 300 ms time lag.

We verified these condition-independent peaks using a permutation based approach which was implemented with the Mass Univariate ERP Toolbox (Groppe, Urbach, & Kutas, 2011).First, the TRF of each participant, condition, and channel was baseline corrected within the function *sets2GND* using time lags from −100 to 0 ms. Second, two-sided t-values were calculated within the function *tmaxGND* using a time-window from 0 to 450ms time lag.Finally, a time-window was identified as significant when t-values exceeded a significant threshold of p <0.05.

Within the significant time-windows we determined individual TRF peaks. For this, we first calculated the standard deviation over channels to derive the global field power (GFP) of the TRF. The GFP indicates the magnitude of a signal across channels. Thereby, it accounts for individual differences in spatial distribution and avoids channel selection (Murray, Brunet, & Michel, 2008). The resulting GFP of each condition were averaged. Next, we searched for the condition-averaged, maximum GFP value in each significant time-window and for each participant. Then, we calculated for each participant the full width at half maximum with respect to the peak to determine individual time-windows. Finally, we averaged over the individual time-windows of the GFP of each condition. This resulted in an average GFP value for each participant, condition, and significant time-window.

### 2.5 Statistics

#### 2.5.1 Preregistered Analyses

Condition differences were analyzed using a linear mixed model (LMM). The analysis was performed in RStudio (version 2021.09.0) using the R package lmer4 (version 1.1-23). For all analyses a categorical fixed factor ‘condition’ with two categories was used, i.e., narrow and wide which were coded 0 and 1, respectively.

For the ERP analyses the response amplitude was predicted for each trial. Participant and channel were included as random factors (Volpert-Esmond, Page-Gould, & Bartholow, 2021):

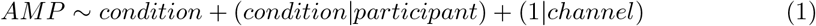

For the ERP model of the beep we ran into singularity issues. The random factor for channel showed a variance of 0 indicating over-specification of this random factor (Volpert-Esmond et al., 2021). We therefore excluded this factor when computing the model for the beep.

For the TRF analyses GFP differences were predicted for individual time averaged peaks. Participants were included as a random factor:

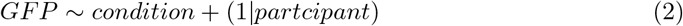

LMM analyses allow the investigation of the random factors participant and channel. For this, the intraclass correlation coefficient (ICC) was used which represents the amount of variance in the predicted variable that is explained by the random factors (Lorah, 2018; Volpert-Esmond et al., 2021). Variances for each factor were calculated using an intercept only model for the analyses of ERPs (*AMP* ∼ 1 + (1 |*partcipant*) + (1 |*channel*)) and TRFs (*GFP* ∼ 1 + (1 |*partcipant*)). ICCs were calculated by dividing the variance of participant or channel by the total variance.

Fixed effects were evaluated using Satterthwaite approximations within the R package lmerTest, which estimates the degrees of freedom to calculate two-tailed p-values. Evidence for an effect were assumed for p-values below .05. We also report standard errors (SE) and 95% confidence intervals (CI).

#### 2.5.2 Exploratory Analyses

We explored reaction time and hit rate in response to the targets, i.e., the target in the narrow condition was the alarm and in the wide condition the beep. For this we used all trial-level responses, i.e., also those that were not considered in the ERP-analyses.

Individual reaction times between conditions were compared using a generalized linear mixed model (GLMM) with an inverse Gaussian distribution to account for a positive skew in reaction time data (*Reactiontime* ∼ *condition* + (*condition*|*partcipant*)).

Hit rate followed a binomial distribution, as hits and misses were coded one and zero, respectively. Differences between conditions was therefore compared using a GLMM with a binomial distribution and logit link function (*Hitrate* ∼ *condition*+(*condition*|*partcipant*)). The statistical significance of differences in reaction times or hit rate between the alarm in the narrow and the beep in the wide condition was evaluated using the Wald Chi-square test.

## 3 Results

### 3.1 Behavioral responses

Figure 2 shows the reaction times and hit rates in response to targets. In other words, the response to the alarm in the narrow condition is compared to the response to the beep in the wide condition. Estimated mean reaction times in response to the alarm in the narrow condition were 0.814 seconds (b=1.22; SE=0.064) and to the beep in the wide condition 0.81 seconds. Reaction times did not differ between the two targets (Figure 2A; b=-0.008; SE=0.045; p=0.869). However, the chance of hitting a target in the narrow condition was 96.2% and in the wide condition 68%. The beep was significantly less often detected than the alarm (Figure 2B; b=-2.478; SE=0.344; p*<*0.001).

**Figure 2:**
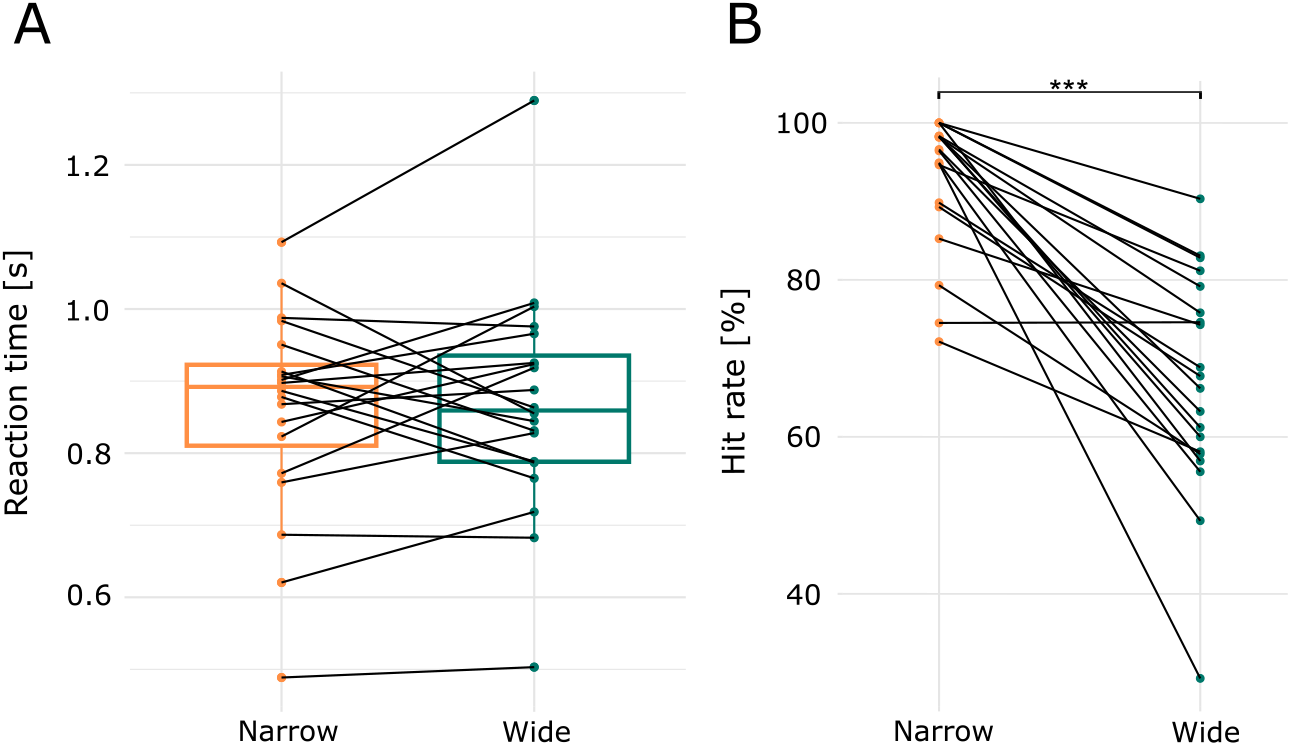
Reaction time (**A**) and hit rate (**B**) in response to the targets of the respective condition, i.e., the alarm and the beep in the narrow and wide condition, respectively. Individual averaged reaction times and hit rates are connected by the black lines. ***p*<*.001

### 3.2 Event-related potentials

We investigated ERPs in response to task-relevant and irrelevant sounds. The alarm was relevant in the narrow and the beep relevant in the wide condition. For these sounds we investigated the P3 response, and expected that targets show larger P3 amplitudes than non-targets. The irrelevant sounds were ignored in both conditions. Here we investigated the N1 response and expected a lower amplitude in the wide compared to the narrow condition.

#### The alarm

Figure 3A shows the grand average ERP (i.e., averaged over participants and selected channel) in response to the alarm in the two conditions. We see a clear N1 peak around 100 ms, a P2 peak around 200 ms, and a P3 response that starts around 300 ms. The topographies of the narrow condition shows a typical parietal P3 distribution (Polich, 2007). The mean amplitude of the P3 response for the alarm in the narrow condition was 4.2 *μ*V with a significant mean amplitude decrease in the wide condition of −2.3 *μ*V (SE=0.53; CI=[−3.39, −1.27]; p*<*.001).

**Figure 3:**
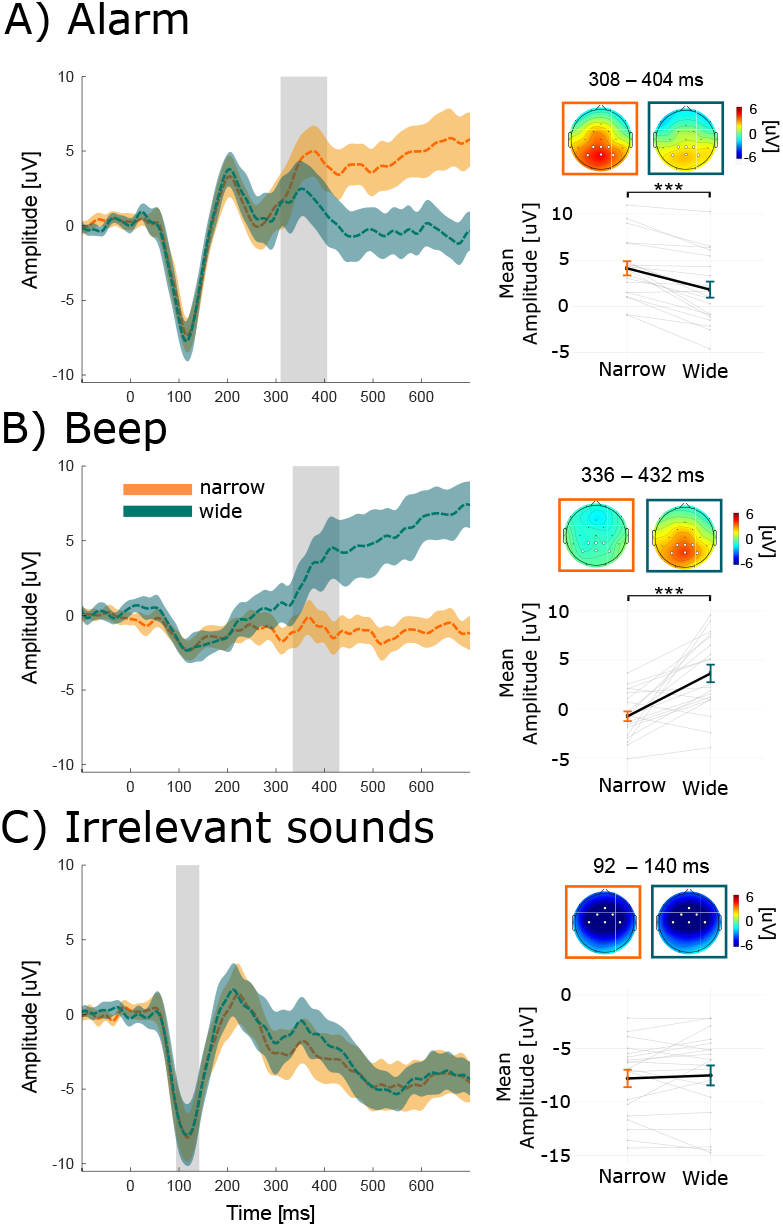
ERPs of each sound and condition, averaged over participants and selected channels. The selected channels are marked in white in the topographies. The narrow and wide condition are marked with orange and green, respectively. Color shades indicate the 95% confidence interval. The gray area indicates the average time-window and the topographies show the average amplitude over this time-window. Note, that individual time-windows were used for the statistical comparison. Below the topographies, the fixed effects (thick black lines) and the variability of effect between individuals (i.e., each grey line corresponds to one participant) are displayed. *** p*<*.001

Computing the ICC showed that variance between people and channel accounted for 12.2% and 0.1% of the total variance, respectively (see supplement table S2 for the results of the random effect models).

#### The beep

Figure 3B shows the grand average ERP in response to the beep. For the beep we neither see a clear N1, nor a P2 peak, but a P3 response that starts around 300 ms. The topography also reflects a P3 response to the beep in the wide condition. The mean amplitude of the P3 response for the beep in the narrow condition was -.609 *μ*V with a significant mean increase in the wide condition of 4.1 *μ*V (SE=0.92; CI=[2.58, 6.16]; p*<*.001). The ICC showed that the variance between people and channel accounted for 6% and 0% of the total variance, respectively.

#### Irrelevant sounds

Figure 3C shows the grand average ERP in response to the irrelevant sounds. We see a clear N1 peak around 100 ms and a P2 peak around 200 ms. The mean amplitude of the N1 response for the irrelevant sound in the narrow condition was −7.81 *μ*V which did not differ from the wide condition (b=-0.28; SE=0.48; CI=[-1.02, 0.87]; p=0.558). The topography reflect a frontal N1 response in both conditions. The ICC showed that variance between people and channel accounted for 14.8% and 0.2% of the total variance, respectively.

#### Summary of ERP results

We found evidence for two of our three hypotheses. The alarm and beep both showed a P3 response when they were the target. This shows that participants were able to detect the sounds. The P3 response subsides after approximately 1.5 seconds (see supplementary figure S4). The response to the irrelevant sound did not change. In the following we look at the processing of the entire soundscape.

### 3.3 Processing of the soundscape as a whole

Figure 4A shows the TRF for both conditions (coloured) and condition-independent (black). Condition specific and independent TRFs show a typical shape (Crosse et al., 2016). The gray areas indicate the three time-windows (0-68 ms; 96-192 ms; 216-448 ms time lag) that significantly differed from zero. The topographies show TRF values averaged over the significant time-window. All time-windows show the largest values across the fronto-central channels. For the first and last time-window the values were negative, while the second time-window showed positive values. Figure 4B illustrates the grand average GFP over all participants. In each significant time-window we determined individual time-windows of the GFP and calculated the average amplitude over the individual time-window. The results are shown in Figure 4C. The individual GFP in the third time-window was on average 5.57 in the narrow condition which significantly increased in the wide condition to 6.43 (b=0.77; SE=0.3; CI=[0.15 1.38]; p=0.0211). We did not find significant differences for the first (b=-0.26; SE=0.68; CI=[-1.63 1.1]; p=0.705) and second (b=0.11; SE=0.59; CI=[-1.07 1.3]; p=0.853) time-window.

**Figure 4:**
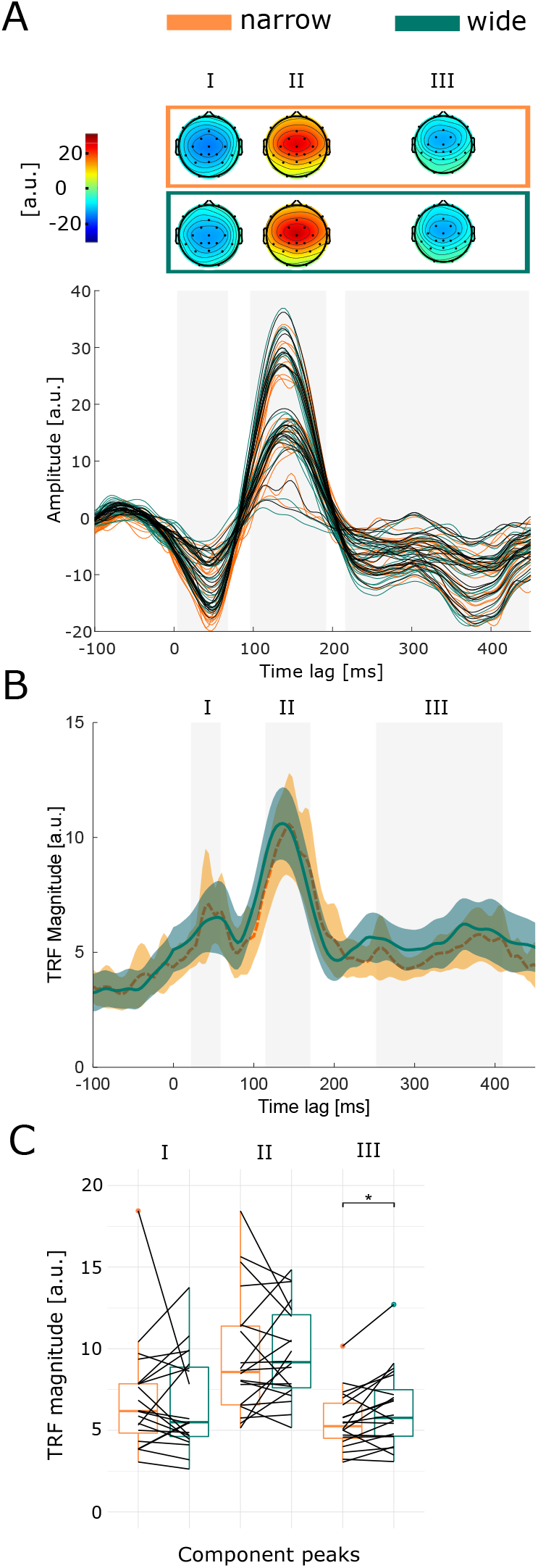
**A)** The topographies show the significant time-windows averaged over time. Below are the TRFs for each channel, averaged over participants. Orange and green mark the narrow and wide condition, respectively. Black marks the TRF calculated over both conditions. The significant time-windows are marked in grey. **B)** The GFP of the TRF is shown for each condition, averaged over participants. Colored area mark the 95% confidence interval. For each participant an individual time-window within the significant time-window was calculated. The grey area marks the average individual time-window. **C)** Boxplots show the differences of the individual time-window for each significant time-window. Each line represents one participant. * p*<*.05

The ICC of the third time-window showed that variance between people accounted for 73.3 % of the total variance indicating a large between-person variance. Note, that the high between-person variance of the TRF compared to the ERPs is the result of using averaged compared to trial-level data, respectively.

One participant showed extremely high standard deviations (see supplement figure S8) across all channel, thus we excluded this participant and ran the analyses again, however, this did not change the results (First: b=0.28; SE=0.43; CI=[-0.58 1.15]; p=0.5174; Second: b=0.44; SE=0.52; CI=[-0.61 1.48]; p=0.414; Third: b=0.84; SE=0.31; CI=[0.22 1.47]; p=0.014).From figure 4B and the supplementary figure S8 it appears that the time-windows vary largely between participants, especially with regards to the last time-window. Therefore, we re-analyzed the data by averaging over the significant time-window that are seen in figure 4A. Using the same time-window for each participant, we receive the same results (First: b=-0.03; SE=0.46; CI=[-0.96 0.9]; p=0.95; Second: b=0.13; SE=0.48; CI=[-0.84 1.1]; p=0.791; Third: b=0.628; SE=0.26; CI=[0.106 1.15]; p=0.026)

## 4 Discussion

We investigated auditory attention using a mobile EEG setup while participants completed a complex audio-visual-motor task with a rich soundscape. We manipulated auditory attention while keeping the complexity of the soundscape constant. In both conditions participants attended to a target. In one condition, this target was a clearly audible alarm originating from one direction which required a narrow attentional focus. In the other condition, this target was a beep originating from different directions which required attention to the whole soundscape, i.e., a wide attentional focus.

Behaviourally we found, that the sound that was assumed to be more difficult to detect (i.e. the beep), was indeed less often detected than the sound that was assumed to be easy to detect (i.e., the alarm). We found a larger P3 ERP response if a sound was a target compared to the same sound if it was not the target. In other words, the response was larger for the alarm in the narrow compared to wide condition and for the beep in the wide compared to narrow condition. We observed the difference around 300 ms after stimulus onset. Contrary to our expectation, we did not find a clear difference in the N1 ERP response to stimuli that were irrelevant in both conditions. We also found a significantly larger TRF response in the last time-window of the TRF for the wide compared to narrow condition.

### Processing of relevant stimuli

The observed P3 response for the targets indicates that participants correctly identified the target. The P3 response is related to attentional processes (Polich, 2007) and our findings show the robustness of this attention effect even in a complex task with visual input, auditory instructions, and motor responses. Thereby it lines up with beyond-the-lab studies that showed the P3 during walking (Debener, Minow, Emkes, Gandras, & de Vos, 2012), biking (Scanlon et al., 2019), driving Protzak and Gramann (2018), and office work (Hölle, Meekes, & Bleichner, 2021). Importantly, the P3 morphology was comparable between conditions (also when looking at the individual participant data in supplementary figure S5 and S6), despite the fact that the targets differed in their characteristics. The alarm was louder than the beep, it came always from the same direction, and was the only sound coming from that direction. The beep, originated from different directions and was embedded into the other sound streams. The behavioral results show that the alarm was easier to detect than the beep. The reaction time for the detected sounds was not significantly different. This shows that a sound which is hard to detect, i.e., acoustically not salient, can elicit a clear attention response if it is considered task-relevant.

### Processing of irrelevant stimuli

Regarding the irrelevant sounds we did not find a difference between conditions in the N1 response. We expected that attention to the beep (i.e., the wide condition) would draw attention to the whole soundscape and in turn also lead to a stronger processing of the irrelevant sounds. This manipulation was apparently not strong enough to produce a difference in the N1 component.

Nevertheless, we can draw conclusion from the observed ERP morphology. The alarm and irrelevant sound elicited an N1 with a clear peak and strong deflection (∼7-8 *μ*V), while the beep elicited an N1 that was smaller (∼ 2-3 *μ*V) and smeared. We interpret the clear peak of the alarm and irrelevant sounds as an indication that these sounds were easily detectable (i.e., acoustically salient) compared to the beep as the N1 is sensitive to sound intensity (Näätänen, 1982; Näätänen & Picton, 1987). Furthermore, early auditory responses indicate awareness of a stimulus (Schlossmacher, Dellert, Bruchmann, & Straube, 2021). Thus, the clear peak of the alarm and irrelevant sounds might indicate that these sounds showed a different early processing compared to the beep.

### Processing of the soundscape as a whole

We found reliably TRFs in response to the complex soundscape (including language and non-language stimuli) in this complex task, with three time-windows which significantly differed from zero. These time-windows have repeatedly been reported for speech and music stimuli (e.g., Hausfeld, Riecke, Valente, & Formisano, 2018; Horton, D’Zmura, & Srinivasan, 2013; O’Sullivan et al., 2015), however not for other complex soundscapes.

We further expected a difference in processing of the whole soundscape between the two conditions. We used the beep in the wide condition to implicitly direct the participants attention towards the whole soundscape. In the narrow condition most of the soundscape could be ignored. We found a significant but small increase of processing in the wide condition after controlling that the effect was not due to the targets. Interestingly, the difference appeared in the last time-window. When tracking the response to an attended and continuous speech stream, an enhanced responses in late time-windows is observed compared to an ignored stream (Holtze, 2021; Horton et al., 2013; Jaeger et al., 2020; Kong, Mullangi, & Ding, 2014; Mirkovic et al., 2019; Petersen, Wöstmann, Obleser, & Lunner, 2017). As we expected that participants attend the whole sound scape more in the wide compared to narrow condition, it is plausible that we observed a difference in the last time-window.

There are several reasons why the observed difference was small. On the one hand, participants did not attend the whole soundscape much more in the wide than in the narrow condition. This would also explain the low hit rate for the beep. On the other hand, there was no incentive to ignore the soundscape in the narrow condition, which might have increased the response to the soundscape in the narrow condition. We conclude that our results are an indication that differences in the processing of the whole soundscape are found in late time-windows.

### Random effects of the models

We further investigated the random effect structure for a better understanding of the variance that contributed to our models (Lorah, 2018; Volpert-Esmond et al., 2021). Interestingly, the between-person ICC of the response to the alarm was twice as large compared to the beep. This indicates that in naturalistic soundscapes, reliable sounds (such as the alarm which was presented from the same direction with the same sound intensity) produce a more reliable trial-level response than unreliable sounds (such as the beep which was presented from different directions and with different sound intensities).

The low between-channel variance indicates that we used channels that were related to the investigated components (i.e., N1 and P3; Volpert-Esmond, Merkle, Levsen, Ito, & Bartholow, 2018)). Furthermore, the selected channels had a close proximity. For the beep we even had to exclude channel as a factor, as we ran into singularity issues.

### Translation to the operating room

We designed our study to contain several factors that characterize the working environment in an operating room, i.e., multiple sound streams from different locations with relevant and irrelevant sounds, speech and non-speech sounds, and a visual-motor task. Our results demonstrate that it is feasible to study auditory attention in such a complex scenario. We observed a clear N1 peak for sounds that were acoustically salient, a P3 response for relevant sounds, and a TRF response for the whole soundscape. Thereby, our study is a step towards studying auditory responses in the operating room using mobile EEG.

Eventually, we want to know how the individual perceives the soundscape in the operating room and when sounds become a burden. Therefore, one must be aware of the challenges of studying sound processing in the operating room: First, the soundscape of an operating room is an uncontrolled setting in which the presentation of sounds is ethically not viable. Therefore, it is necessary to relate the natural soundscape to the EEG recording. We showed that meaningful EEG responses can be measured in a complex soundscape. In a following step, we suggest to use smartphone-based technology that enables the simultaneous recording of EEG and audio features in a data protected way (Blum, Hölle, Bleichner, & Debener, 2021; Hölle, Blum, Kissner, Debener, & Bleichner, 2021), and apply it to the operating room. This way, responses to naturally occurring sounds can be measured. Second, surgery staff is exposed to the soundscape for several hours per day. Investigating changes in sound processing over the day requires long-term recordings. Here one could use minimal and unobtrusive EEG set-ups like the cEEGrid (Debener, Emkes, De Vos, & Bleichner, 2015), that can be used to study EEG responses to auditory stimuli (Holtze, Rosenkranz, Jaeger, Debener, & Mirkovic, 2022; Meiser & Bleichner, 2022) over more than six hours (Hölle, Meekes, & Bleichner, 2021). Lastly, the cognitive load (e.g., working memory and attentional capacities) varies during a surgery over time and between staff members which likely affects auditory processing. van Harten et al. (2021) observed that surgery staff with high workload is more likely distracted by irrelevant sounds than surgery staff with low workload. However, this relationship is simplified as high load can also reduce the processing of irrelevant sounds (Brockhoff, Schindler, Bruchmann, & Straube, 2022). Studying the relationship between load and auditory processing in the operating room is therefore necessary to understand the effect that sounds have on the surgery staff.

## Conclusion

We showed that ERPs, as well as TRFs, are useful tools to study different aspects of sound perception in complex sound environments. To balance between high control over stimuli and the uncontrolled operating room we developed a laporatory experiment with a naturalistic soundscape. In this scenario ERPs are robust to detect attention responses to specific sounds while TRFs can measure responses to an uncontrolled soundscape. Our results demonstrate that we can use mobile EEG in a complex acoustic-visual-motor task to study auditory perception and is thereby an important step towards understanding auditory attention in uncontrolled settings.

## Footnotes

1 https://github.com/labstreaminglayer/liblsl-Matlab, v1.14

2 https://github.com/labstreaminglayer/App-Input, v1.15

3 https://github.com/labstreaminglayer/App-AudioCapture, v1.14

4 https://github.com/labstreaminglayer/App-LabRecorder, v1.14

5 filter order = 16500, transition bandwidth = 0.1 Hz, cutoff frequency (−6dB) = 0.05 Hz

6 filter order = 220, transition bandwidth = 7.5 Hz, cutoff frequency (−6dB) = 33.75 Hz

7 filter order = 1650, transition bandwidth = 1 Hz, cutoff frequency (−6dB) = 0.5 Hz

8 filter order = 220, transition bandwidth = 7.5 Hz, cutoff frequency (−6dB) = 33.75 Hz

## Notes

https://doi.org/10.5281/zenodo.7147701

### Competing Interest Statement

The authors have declared no competing interest.

