## SupplementaryMaterial for "Investigating the attentional focus to workplace-related soundscapes in a complex audio-visual-motor task using EEG"

### Supplementary Material

### 1 Deviations from our preregistration

General deviations:

1. We originally referred to the beep as the odd tone.

Deviations from planned preprocessing:

1. We did not specify the ICLabel threshold of .9 to reject components.
2. Regarding the P3, we initially searched for a peak between 250 and 400 ms. However, for some participants this resulted in finding the the P2 and not the P3 time-window. Therefore, we searched for a peak between 300 and 400 ms.
3. Regarding the data epochs for the TRF, we did not consider that the total length might significantly vary between conditions. Thus, epoch exclusion until the conditions have a similar length was added as a preprocessing step.

Deviations from planned analyses:

1. We did not specify during the preregistration how we deal with models that run into singularity issues, but trimming random factors with zero variance is recommended (Volpert-Esmond et al., 2021). Thus, we decided to drop the random factor ”channel” for the ERP analyses of the beep.
2. As proposed in the preregistration, we planed post-hoc power analyses. However, using trial-level data our computational power was not sufficient to calculate a power analysis (i.e., it would have taken several weeks to conduct such analysis). Besides, we noticed that the use of post-hoc power analysis might be misleading, as it diverts from the true power to detect a significant effect (Zhang et al., 2019).

### 2 Supplementary Method

#### 2.1 Tetris Game Design

The Tetris space consisted of 15 layers and each layer covered a 5x5 area. The layers and area formed a grid with equally sized cubes, thus forming a space for 15x5x5 cubes (see Figure 1). A block was formed from several cubes, resulting in blocks of ten different shapes (see Figure 2). A block was moved with the arrow keys or rotated cloackwise with the A,S,and D keys. The blocks dropped at a fixed speed until they were placed at the bottom of the space or on top of other block(s). While a block dropped, it was transparent and only its boarders were visible. If a block was placed, it was colored depending on the layer in which it was placed. Each layer has its own color. Thus, if a block covered more than one layer, its cubes were colored differently. To the right of the Tetris space, the current score, the shape of the next block, and the color code of the layers was shown.

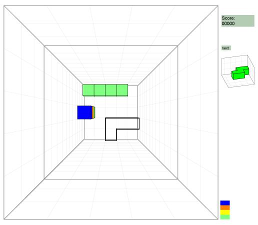

Figure S 1: Tetris Box

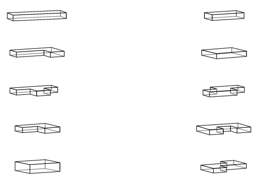

Figure S 2: Tetris Blocks

Within the game, there were three ways how points were granted or subtracted:
First, if a layer was completely covered with cubes, it was deleted and 100 points were granted. If the blocks stacked too high, i.e., if at least one cube exceeded the 13th layer, blocks in all layers were deleted and 100 points subtracted. Thereby, participants had enough time (i.e., 3 layers) to navigate the blocks even if blocks were stacked high. Furthermore, by deleting all blocks, continuity of the game was guaranteed.
Second, participants received instructions to press the space bar when hearing the targets, i.e., the alarm or beep in the narrow or wide condition, respectively. Correct presses granted 50 points, missing a target subtracted 50 points.
Third, following the task-relevant speech granted 100 points while not following it subtracted

100 points. The task-relevant speech gave instructions to place the block in the upper or lower and left or right corner of the 3D space. A corner was defined as a 2x2 area in any layer. If at least one cube of a block was placed in the correct area, the points were granted.

#### 2.2 Auditory Stimulus Material

Three types of auditory stimuli were integrated: a continuous background sound, distinct sound events, and speech. The background sound was adapted from an open-access hospital recording available on youtube [9]. The adaptation was manually done in Audacity® and included the removal of identifiable speech segments and creating chunks of the audio which could be rearranged and duplicated. This way, two versions of the background sound were created which were counterbalanced across conditions.
Regarding the distinct sound events, four sounds were included. Two of them were hospital monitor sounds which served as the irrelevant sounds[10] and one a hospital alarm sound[11]. As the youtube soundfiles contained multiple presentations, one sample of each sound lasting approximately 200 ms was manually extracted with Audacity®. The irrelevant sounds were presented equally often in the experiment. The fourth sound event, the beep, had a frequency of 800 Hz, was about 60 ms long, and was generated in MATLAB.
Two types of speech stimuli were presented. The first type was task-relevant speech. It was generated using a text-to-speech program[12]. The speech instructed participants to place the next block in one of the four corners, thus four instructions were created. Each of the four instructions were presented equally often and the same instruction was never presented subsequently. The second type of speech was task-irrelevant speech. It contained snippets from a German podcast conversation between a male and female speaker. The snippets were manually extracted using Audacity® and presented in a predefined order. Each snippet lasted on average 3.5 (+- 1.5) seconds. Two podcasts were used [13] which were counterbalanced across conditions.
After the initial manual extraction of the sounds, they were further processed in MATLAB. Therefore, each stimulus was matched to the average root-mean-square value of all sounds. The sounds were spatially seperated using the Head Related Impulse function (Kayser et al., 2009), except for the background sound. As the loudness of the sounds varied, they were multiplied by individual factors (i.e., gain) before the spatial separation algorithm was applied. As the beep is presented together with other sounds, five versions of the beep were created. Each version was processed with the same parameters as the respective stimulus. For the beep in the background, no spatial separation was performed. Table 1 provides a detailed overview of the applied parameter for each stimulus.

| Stimulus | Location | Listener Position | Speaker Position | Gain |
|---|---|---|---|---|
| Irrelevant Sounds | Cafeteria | 2 | A | 10 |
| Alarm | Cafeteria | 2 | E | 10 |
| Relevant Speech | Cafeteria | 2 | D | 25 |
| Irrelevant Speech | Cafeteria | 2 | B | 6 |

Table S 1: *HRIR processing parameters of auditory stimuli.*

#### 2.3 Stimulus Presentation

At the start of the game, the 3D-space was empty. A game started after a countdown counted from three to one. A Tetris block always started in the middle of the upper most layer, i.e. the 15th layer. The shape of a block was randomly chosen from one of the ten shapes.
Within the game, auditory stimuli were presented using Psychtoolbox 3 (Kleiner et al., 2007). For each presented stimulus a trigger marker was generated using a Lab Streaming

[9] https://www.youtube.com/watch?v=qR9YzVqO9Zg
[10] https://www.youtube.com/watch?v=4NXe9pwEgN4&list=OLAK5uy_m0wcbXgo3PYKOigdAPPPf0s9y2F7ZXtp0&index=127
[11] https://www.youtube.com/watch?v=95siaTtQR-c&list=OLAK5uy_m0wcbXgo3PYKOigdAPPPf0s9y2F7ZXtp0&index=128
[12] www.notevibes.com(Notevibes, 2021)
[13] https://www.ndr.de/wellenord/Allein-unter-Moerdern-Sabine-Thiesler,kunstmichmal166.html and https://www.ndr.de/wellenord/Beruehren-verboten-Nicht-bei-Julia-Brunner,kunstmichmal150.html

Layer based software [14]. The background sound was presented throughout the game. Each of the four relevant speech instructions were presented 12 times. Each snippet of the irrelevant instructions was presented once in a predefined order. In total 48 irrelevant speech snippets were presented. The two irrelevant sounds were presented 24 times each. The alarm and beep were presented 48 times each. However, the beep could occur together with other stimuli. At the start of the game the beep was randomly positioned in each stimulus. The onsets between the irrelevant sounds/alarm and beep were close, potentially influencing the ERP analysis. Therefore, for each time a sound was presented together with a beep, the sound and beep were again added to the pool of stimuli. This resulted in a variation of sound presentations across conditions and participants. For the ERP analyses only the 48 trials without other interfering sounds were used.

Auditory stimuli were presented randomly depending on the layer of a block: The first auditory stimulus during a block occurred between layer 12 to 6, the second stimulus three to six layers after the first one, and so on. However, there were some restrictions. Task-relevant speech could occur consecutively, but the same speech instruction (i.e., placing the block in the same corner) could not occur consecutively to ensure that the blocks do not stack too high. The beeps did not occur during the first five stimuli. The position of the beep within another stimulus was randomly defined at the beginning of each condition. Thus, it was different for each participant and condition, but fixed for a stimulus. Note, that the beep could occur at a different position for each of the four instructions, two irrelevant sounds, and 48 task-irrelevant speech snippets. The participants were not aware of the presentation frequency or order. They were only informed about the approximate length of a game.

#### 2.4 Training

To get acquainted to the game, participants received written instruction and performed four training games. During the first training, block rotation and placement was trained without any auditory stimuli and lasted ten minutes. During the first half, blocks dropped at a low speed. During the second half, blocks dropped at normal speed.

The second training game introduced the background sound and task-relevant speech, as it was relevant for both conditions, and lasted approximately one minute. Here, participants received visual feedback whether they correctly followed the instructions.

The third and fourth training games included all stimuli and were condition specific and therefore, played before the respective condition. They lasted approximately two minutes each. In the narrow condition training, participants received visual feedback whether they correctly detected the alarm. Prior to this training game the alarm was presented to them. In the wide condition training, participants received visual feedback whether they correctly detected the beep. Prior to this training game the beep was presented to them once alone and once included in the other stimuli.

A game ended after all stimuli were presented. For the games of each condition this was the case after approximately 18 minutes. For the speech training, five instructions were presented. For the condition specific training, each stimulus was presented twice.

[14] `www.github.com/labstreaminglayer/liblsl-Matlab`

### 3 Channel Layout

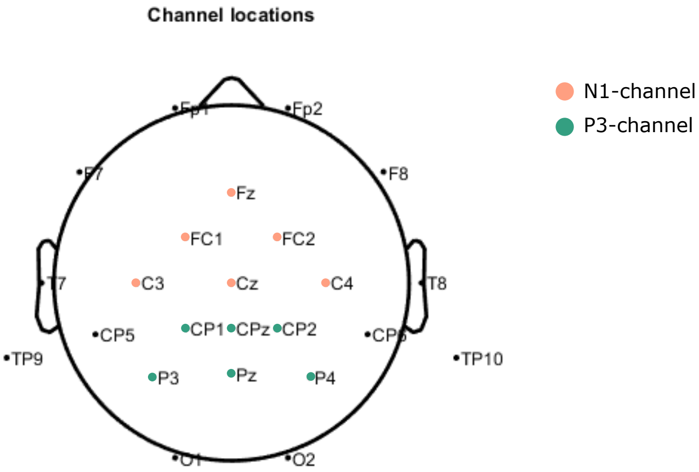


Figure S 3: A mobile 24-channel setup was used for this study. The colours orange and green indicate the channels that were used for the N1 and P3 ERP analysis, respectively.

### 4 Random effect model results

| Stimulus | Group | Variance | Std. Dev |
|---|---|---|---|
| Alarm | Participant | 12.08 | 3.476 |
| | Channel | 0.11 | 0.325 |
| | Residual | 86.71 | 9.312 |
| Beep | Participant | 6.002 | 2.450 |
| | Channel | 0 | 0 |
| | Residual | 94.028 | 9.697 |
| Irrelevant sounds | Participant | 13.44 | 3.6661 |
| | Channel | 0.162 | 0.4026 |
| | Residual | 76.9825 | 8.7740 |
| TRF III | Participant | 3.019 | 1.737 |
| | Residual | 1.099 | 1.049 |

Table S 2: *Random effect results of the intercept model for the alarm, beep, irrelevant sounds, and the third TRF time-window*

### 5 Long time-window of target ERPs

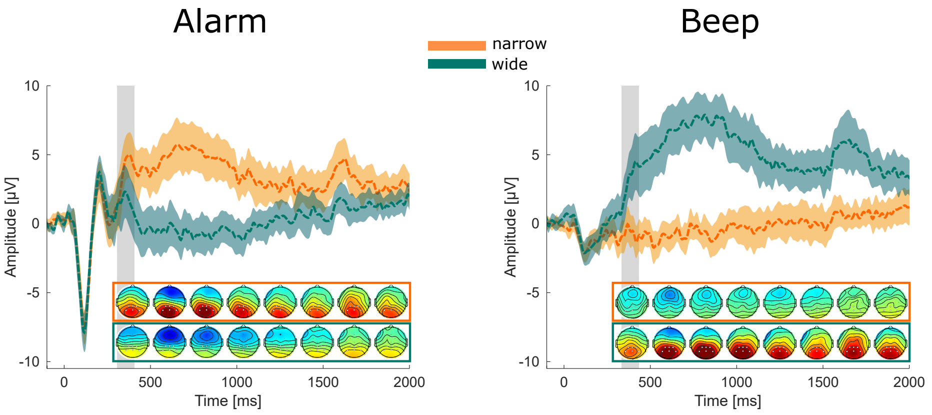


Figure S 4: The ERPs are the same as in figure 3. Topographies show time-windows from 300 to 1900 ms over time in steps of 200 ms.

### 6 Individual participant ERPs

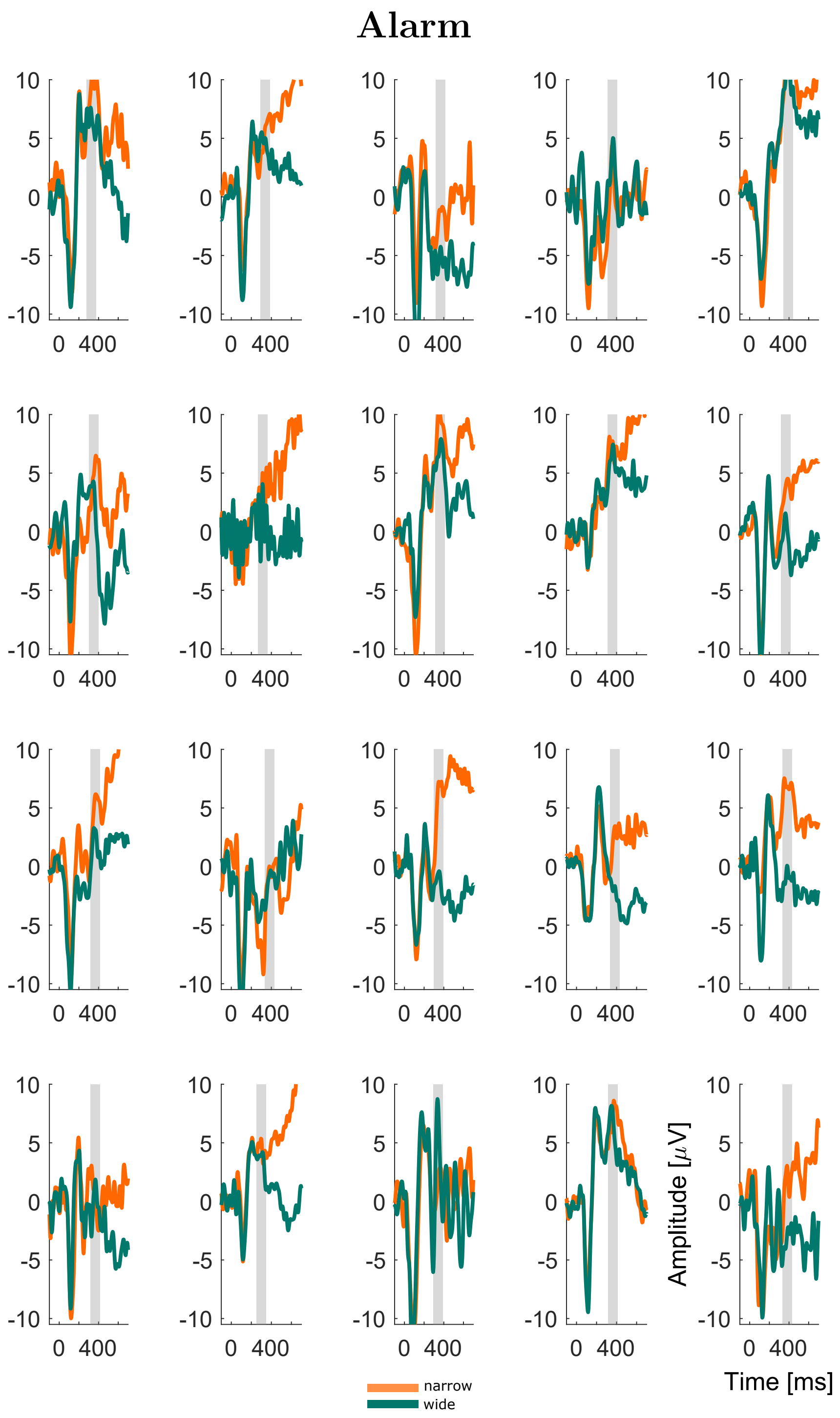


Figure S 5: Individual participant data in response to the alarm averaged over selected P3-channel and trials. Gray area marks the individual time-window used for the statistical comparison.

### Beep

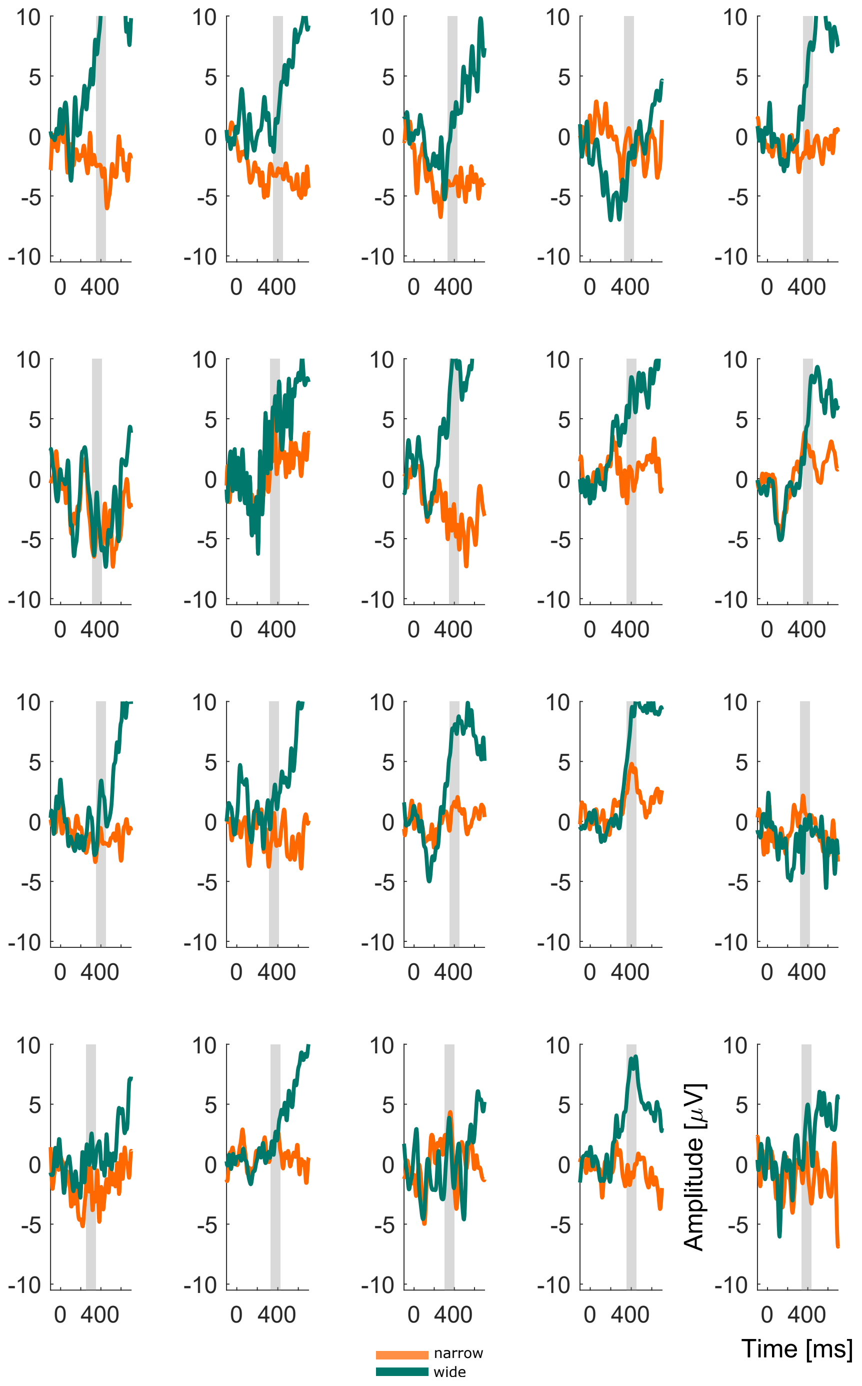


Figure S 6: Individual participant data in response to the beep averaged over selected P3-channel and trials. Gray area marks the individual time-window used for the statistical comparison.

### Irrelevant Sound

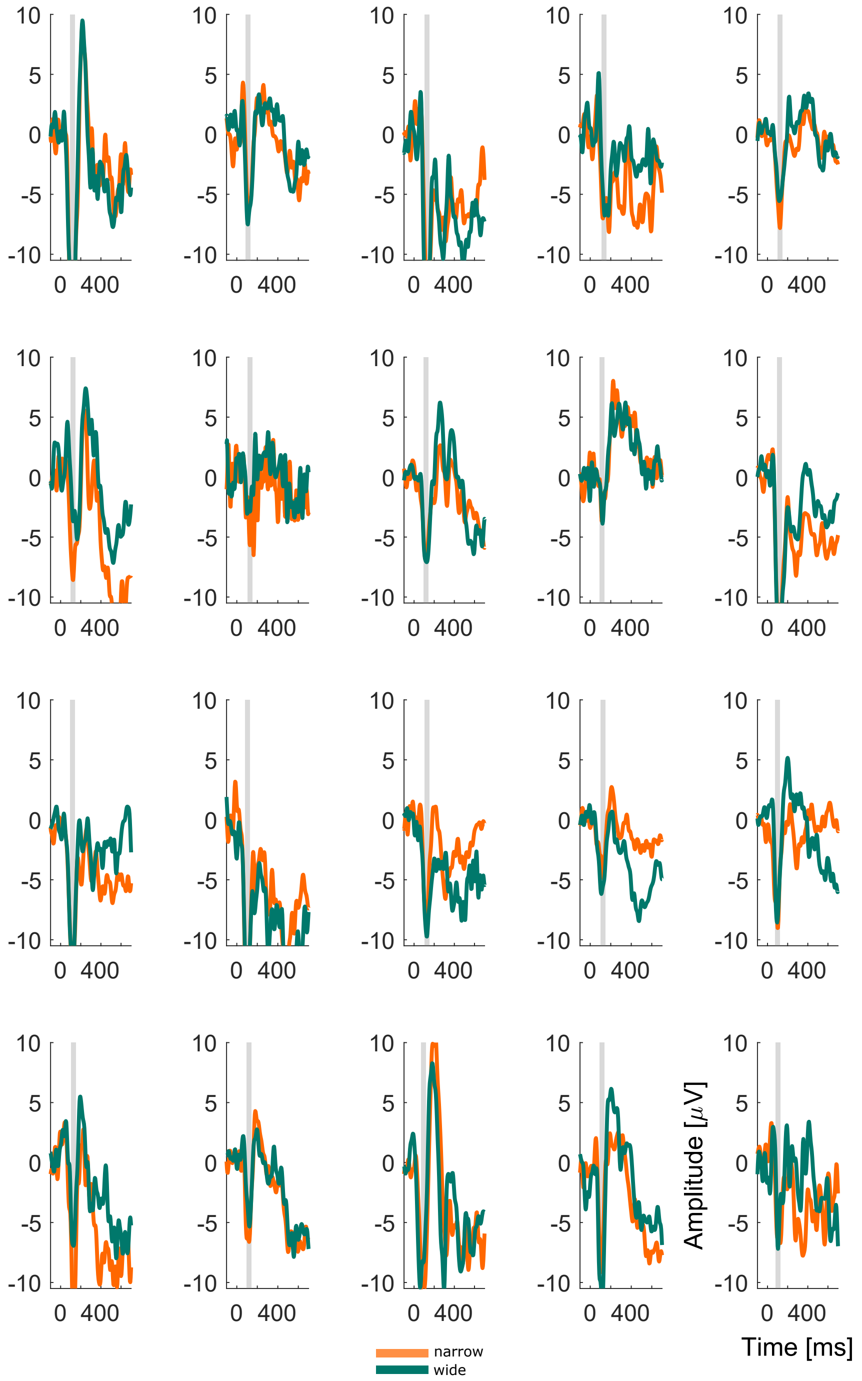


Figure S 7: Individual participant data in response to the irrelevant sounds averaged over selected N1-channel and trials. Gray area marks the individual time-window used for the statistical comparison.

855 # 7 Individual participant TRFs

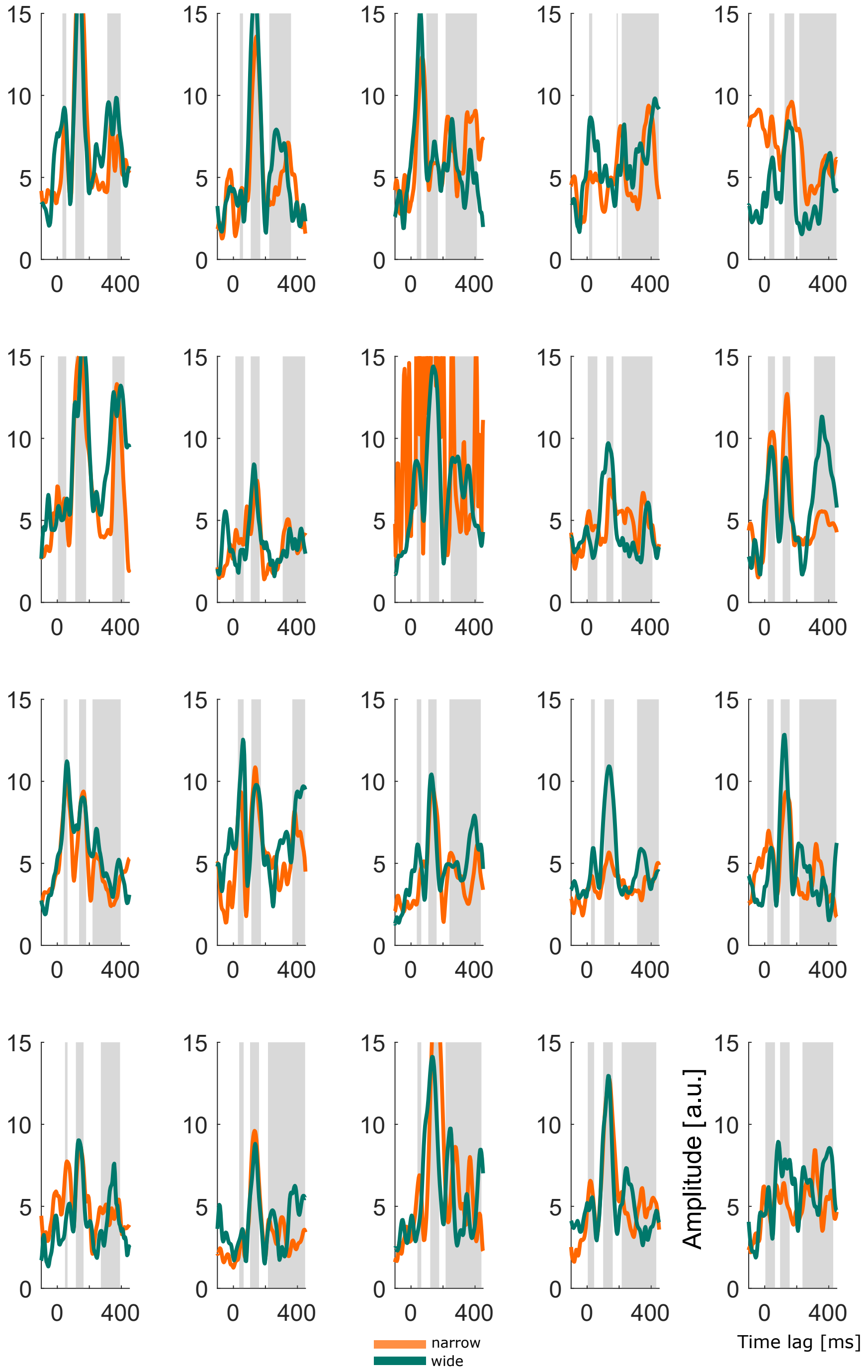


Figure S 8: Globald field power of the TRF for each participant in response to whole soundscape. Gray areas mark the individual time-windows used for the statistical comparison.
